# Mud sticks after all: Chronically implanted microelectrodes but not electrical stimulation cause *c-fos* expression along their trajectory

**DOI:** 10.1101/203877

**Authors:** Patrick Pflüger, Richard C. Pinnell, Nadja Martini, Ulrich G. Hofmann

## Abstract

The goal of CNS implanted devices is to build a stable brain-machine-interface. The brain tissue response to the foreign body limits the functionality and viability of this brain-machine connection. Notably the astrocytic glial scar formation and inflammation with resulting neuronal cell loss is considered to be responsible for the signal deterioration over time. We chronically implanted a polyimide microelectrode in the dorsolateral striatum of rats. First, we analyzed the *c-fos* immunoreactivity following high frequency stimulation (HFS) of the dorsolateral striatum and second, using GFAP and ED1 immunocytochemistry, the brain tissue response. Acute as well as chronic HFS showed no significant change of neuronal *c-fos* expression in the dorsolateral striatum and corresponding cortical areas. We found that the sole chronic implantation of a polyimide microelectrode leads to a reaction of the surrounding neurons, i.e. *c-fos* expression, along the implantation trajectory. We also observed the formation of a glial scar around the microelectrode with a low number of inflammation cells. Histological and statistical analysis of NeuN positive cells showed no ‘kill zone’, which accompanied neuronal cell death around the implantation site.

## INTRODUCTION

Electrical high frequency stimulation (HFS) is an established method to treat advanced Parkinson’s disease (Lozano et al., 2002, Garcia et al., 2005, Gubellini et al., 2009), and is commonly also utilized for the treatment of obsessive compulsive disorders and depression (Mayberg et al., 2005, Greenberg et al., 2010). However, a generally agreed upon understanding of the detailed mechanism of HFS is still missing (Benabid et al., 2009), since somewhat controversial HFS of target structures can cause inhibition (Benazzouz et al., 1995, Maurice et al., 2003) or excitation (Hashimoto et al., 2003, Hammond et al., 2008) depending not only on the stimulated target but its input and output regions as well, reviewed in Feuerstein (Mantovani et al., 2009, Feuerstein et al., 2011) or Udapa and Chen (Udupa and Chen, 2015).

Considerable effort is under way to improve DBS by ameliorating its side effects and achieving a better patient outcome. This ambitious goal may be achieved in the future with a multitude of approaches like contingent (closed-loop) stimulation (Little et al., 2013), non-periodic stimulation patterns (Brocker et al., 2013, Brocker et al., 2017), or by deconstructing current four-ring stimulation electrodes into smaller, more numerous microelectrodes (Martens et al., 2011, Kees et al., 2015). However, it has to be emphasized that by decreasing the dimensions of the clinically used stimulation leads, one reaches out into the realm of microprobes and microstimulation - a field with plenty of challenges of its very own (Turner et al., 1999, Polikov et al., 2005, Biran et al., 2007, Leach et al., 2010).

Different cells are involved in inflammation processes and wound healing after implanting a device into the CNS (Polikov et al., 2005, Biran et al., 2007). Microglia as a ‘*first responder*’ is the main cellular player of the acute response to the changed environment (Davalos et al., 2005). They also contribute during the chronic response with the formation of the glial scar (Röhl et al., 2007). Astrocytes are mainly responsible for the reactive gliosis process (Reier, 1986). The viability of the surrounding neurons and their location with respect to the electrode, are crucial for a stable signal transmission to occur (Biran et al., 2005, Polikov et al., 2005). Due to the violation of both the integrity of the CNS by implanting an electrode, and the following foreign body reaction, the brain-machine connection deteriorates. In particular, in chronically implanted electrodes the glial scar formation and neuronal cell loss (‘kill zone’) are held responsible for an overall signal degradation (Edell et al., 1992, Liu et al., 1999, Biran et al., 2005).

Numerous approaches were made for the purpose of improving the brain-machine connection and alleviating the brain tissue response, including: the electrode design (Hofmann et al., 2006) and material (Csicsvari et al., 2003, Kipke et al., 2003), the coating of the shank (Ludwig et al., 2006, He et al., 2007) and the implantation techniques (Kim et al., 2004, Wise et al., 2004) for minimizing the tissue damage and foreign body response (Leach et al., 2010).

To this end we conducted a study to examine the effects of HFS in the dorsolateral striatum by microelectrodes, and the brain tissue response, in a rodent model for up to 10 weeks. The targeted brain area features somatotopically organized corticostriatal connections (Voorn et al., 2004), and already served as model region to illuminate the neurochemical effects of DBS (Hiller et al., 2007).

In the current study, post-mortem immunochemistry was used to probe neuronal (NeuN as neuronal marker) activation by expression of *c-fos* (Faull, 1989, Bullitt, 1990, Herrera and Robertson, 1996, Wilson et al., 2002), astrocytic activity by glial fibrillary acidic protein (GFAP), and microglia activity by anti-CD68 (ED1), in order to determine the inflammatory reaction to the chronic implantation of the microelectrode (Turner et al., 1999, Grill et al., 2009, McConnell et al., 2009a, Beck et al., 2010).

## MATERIAL AND METHODS

### Ethics statement

All procedures involving animals and their care were conducted in conformity with relevant institutional guidelines in compliance with the guidelines of the German Council on Animal Protection. Protocols were approved by the Animal Care Committee of the University of Freiburg under supervision of the Regierungspräsidium Freiburg (approval G13/97) in accordance with the guidelines of the European Union Directive 2010/63/UE.

### Animals

Female Sprague-Dawley rats (290-330g; n = 15) were anaesthesized with oxygen (0.15 l/min) and isoflurane (Abbvie, USA); the latter of which was initially set to 4% and gradually lowered to 1.5% after placing the animal into the stereotaxic frame (David Kopf, USA). Animal breathing, reflexes and level of anaesthesia were monitored throughout the duration of the surgery.

Animals were housed after implantation in pairs (Pinnell et al., 2016) under standard lighting (12h light-dark cycle), 22°C and 40% humidity. Animals were allowed access to food and water *ad libitum*.

Chronically implanted rats were sacrificed with an overdose of isoflurane and perfused transcardially with 4% paraformaldehyde (PFA) in phosphate buffer. Their brains were removed, postfixed in PFA for 7 days and stored in 30% sucrose until cutting them in coronal sections (20 μm) along the probe’s implantation trajectory with a cryostat. The sections were collected on glass and stored at −20°C until further processing.

### Electrode implantation

Prior to surgery, all rats underwent several days of handling in order to familiarize them with the experimenter.

Anaesthetized animals were implanted with a single, stiffened, 16-contact polyimide micro probe (IMTEK; Freiburg University) in the left dorsolateral striatum (AP: +0.4, ML: +3.6; from Bregma, DV: −3.7 from dura mater) (Paxinos et al., 1985) after midline incision and reflection of shaved scalp following the description of (Pinnell et al. 2016). The microelectrode was superglued to a 125 μm glass rod prior to implantation to provide accurate positioning and increase rigidity (Richter et al., 2013). Animals were allowed to recover for 13-15 days. The microelectrode was implanted for a mean time of 70 days.

### Stimulation

The stimulation pulse frequency was set to 130 Hz, and each pulse consisted of rectangular, biphasic pulses with a pulse width of 60 μs/phase, a current of 400 μA, and a duration of 5 minutes (AlphaLab SNR System, Alpha Omega GmbH, Germany). The geometrical area of the stimulating Iridiumoxide microelectrode (Mottaghi et al. 2015) was 2500 μm^2^, yielding a charge per stimulating phase of 24 nC/ph and a total stimulating charge of 960 μC/cm^2^. Rats were either stimulated over 6 consecutive days (chronic STIM) or not (chronic SHAM). Following this period, rats were sub-divided into two additional groups, on the basis of them receiving either stimulation or sham-stimulation, 3 hours prior to being perfused with a ketamine/xylene anesthesia mixture. The resulting groups were: Group1 (n=4): Acute STIM / Chronic STIM; Group 2 (n=3): Acute SHAM / Chronic SHAM; Group 3 (n=5): Acute STIM / Chronic SHAM; Group 4 (n=3): Acute SHAM / Chronic SHAM.

**Fig. 1:**
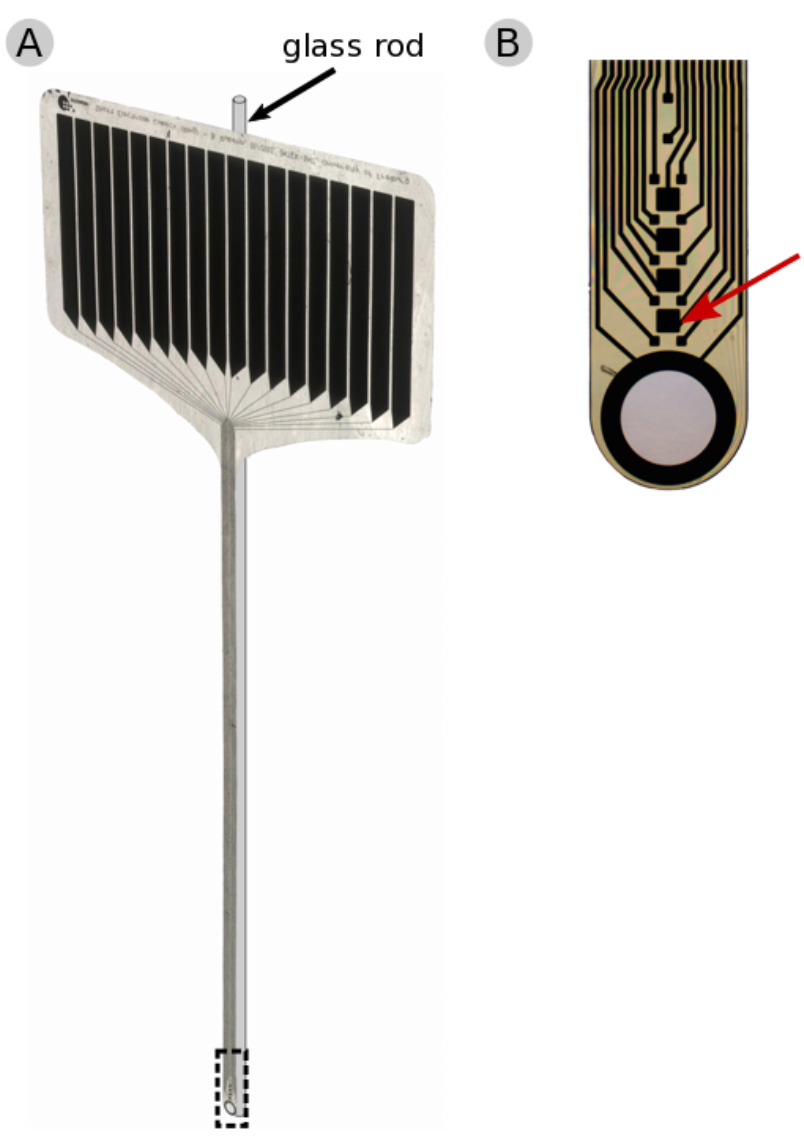
**(A)** 16-contact polyimide micro probe (IMTEK; Freiburg University) with a supporting glass rod. **(B)** Magnified tip of the micro probe, with a red arrow pointing at a stimulation contact (50 μm × 50 μm).

### *c-fos* and NeuN immunofluorescence staining

*C-fos* immunoreactivity was visualized using a double-label immunofluorescent stain for *c-fos* and neuronal nuclei (NeuN). Coronal brain sections were processed and incubated overnight with a polyclonal rabbit anti-*c-fos* antibody (sc-52, Santa Cruz Biotechnology, CA, USA, diluted 1:100). After rinsing in phosphate-buffered saline (PBS), sections were incubated with a fluorescent donkey anti-rabbit IgG conjugated with Alexa fluro 647 (Abcam, CA, USA, diluted 1:1000). For co-visualizing NeuN immunoreactivity - and thus demonstrating *c-fos* within the neuron - sections were rinsed again in PBS, blocked with 10% normal donkey serum (NDS), and incubated for 3 hours with a polyclonal mouse anti-NeuN antibody (Anti-NeuN, Millipore Cooperation, MA, USA, diluted 1:100) (Mullen et al., 1992). After rinsing in phosphate-buffered saline, sections were incubated with a fluorescent donkey anti-mouse IgG conjugated with Alexa fluro 488 (Abcam, CA, USA, diluted 1:1000). Finally, sections were rinsed again in PBS, mounted with DAPI-Fluoromount G (SouthernBiotechnology Associates, Inc., AL, USA) and stored at 4ºC.

### Counting of *c-fos*/NeuN positive cells

For quantitative analysis, 6 sections of each animal were used for counting the *c-fos*/NeuN+ cells along the implantation trajectory. Images of stained sections were taken using a Zeiss microscope equipped with a ProgRes camera, and ProgRes CapturePro 2.7 software (Carl Zeiss, Germany, Jenoptik, Germany). We created composites of the coronal sections using the ImageJ plugin ‘stitching’ (Preibisch et al., 2009), and adjusted the contrast and brightness with the help of Adobe PhotoShop CS6. With the ImageJ software ‘cellcounter’, we first quantified the number of NeuN+ cells per box (100 μm × 100 μm), and in the next step from the colocalized NeuN und *c-fos* sections, the number of *c-fos*/NeuN+ cells. The mean counts of the ipsilateral (stimulated) sides of the coronal sections were compared between groups. For statistical analysis of the NeuN+ cells, we compared the means of the region from 0-100 μm with the background within one group. Therefore, we calculated the difference between those two means and in the next step, the 95% CI. If the 95% CI involved 0 or was >0, we concluded the number of NeuN+ cells in the region immediately surrounding the electrode not to be significantly lower than the background neuronal density (p<0.05).

For the statistical analysis of the *c-fos*/NeuN+ cells we compared, parallel to the analysis of the NeuN+ cells, the means of the region from 0-100 μm with the background within one group and following between the different groups. For the first analysis we calculated the difference between the means of region from 0-100 μm and the background, and in the next step the 95% CI. If the 95% CI did not include 0 or was >0, we concluded the number of *c-fos*/NeuN+ cells in the region immediately surrounding the electrode to be significantly higher than the background (p<0.05). For a secondary comparison between groups we used the software JMP (JMP 13.1.0, SAS Institute Inc., SAS Campus Drive, Cary, NC, USA) and applied a one-way ANOVA and the Scheffé method for post-hoc testing. We defined a level of p<0.05 as statistical significant.

### GFAP and ED1 immunofluorescence staining

To visualize the glial cell and microglial response, we also performed a double-label immunofluorescence staining for glial fibrillary acidic protein (GFAP) and anti-CD68 (ED1). The coronal brain sections were processed and incubated for 3 hours with polyclonal mouse anti-rat-CD68-antibody (AbD Serotec, UK, diluted 1:100). After rinsing in phosphate-buffered saline (PBS), sections were incubated with a fluorescent donkey anti-mouse IgG conjugated with Alexa fluro 488 (Abcam, CA, USA, diluted 1:1000). For visualizing GFAP immunoreactivity, sections were rinsed again in PBS, blocked with 10% normal donkey serum (NDS) and incubated for 3 hours with a polyclonal rabbit anti-GFAP antibody (GFAP, Millipore Cooperation, MA, USA, diluted 1:1000). After rinsing in PBS, sections were incubated with a fluorescent donkey anti-rabbit IgG conjugated with alexa fluro 647 (Abcam, CA, USA, diluted 1:1000). Finally, sections were rinsed again in PBS, mounted with DAPI-Fluoromount G (SouthernBiotechnology Associates, Inc., AL, USA) and stored at 4°C.

### GFAP and ED1 analysis

Four coronal sections along the trajectory of each animal were used to quantify the glial fibrillary acidic protein (GFAP)- and ED1-immunoreactivity. Images of stained sections were taken using a Zeiss microscope equipped with a ProgRes camera with ProgRes CapturePro 2.7 software (Carl Zeiss, Germany, Jenoptik, Germany). Due to the low microglial response to the chronic implantation of the microelectrode, we could not apply a numerical analysis, and a representative picture is shown as an example (Fig. 9). For quantifying the GFAP-immunoreactivity we used ImageJ ‘PlotProfile’ and collected several profiles for each region (cortex, corpus callosum, striatum), separated in medial and lateral, of one section. We calculated the means of one region and site, and subtracted the background immunofluorescence intensities from far away (=Background-corrected immunofluorescence intensity). The profiles of the background-corrected immunoflourescence intensities of the different groups are shown in Fig. 7 A. Furthermore, we calculated the full widths at half maximum (FWHM) to quantify the expansion of the glial scar. For statistical analysis we compared the FWHM between groups. We applied One-way ANOVA and following significant ANOVA, the Scheffé method for post-hoc testing using the software JMP (JMP 13.1.0, SAS Institute Inc., SAS Campus Drive, Cary, NC, USA). We defined a level of p < 0.05 as statistical significant.

## RESULTS

### Effects of the chronic implantation of a microelectrode on *c-fos* expression

All animal groups, independent of their stimulation paradigm, had displayed co-localized *c-fos*/NeuN+ cells along the former location of the microelectrode. Fig. 2 depicts the colocalization of the *c-fos*-labeled cells to NeuN-labeled neurons (white arrows). Statistical analysis of *c-fos*/NeuN+ cells in the region of 0-100 μm from the microelectrode, in comparison to region of 400-500 μm, showed for group 1 a 95% CI of [12.43;31.46], group 2 [38.24;66.21], group 3 [45.27;77.61] und group 4 [27.63;66.93]. Thus, in all 4 groups the number of *c-fos*/NeuN+ cells in the vicinity of the microelectrode is significantly higher than that of the background (p < 0.05). This result demonstrates that a sole chronic implantation of a stiff microelectrode for 10 weeks can cause a *c-fos* expression in neurons on the ipsilateral side (no *c-fos*/NeuN+ cells contralaterally, as there is no implant). The distribution of the *c-fos*/NeuN+ cells along the microelectrode is displayed in Fig. 3.

**Fig. 2:**
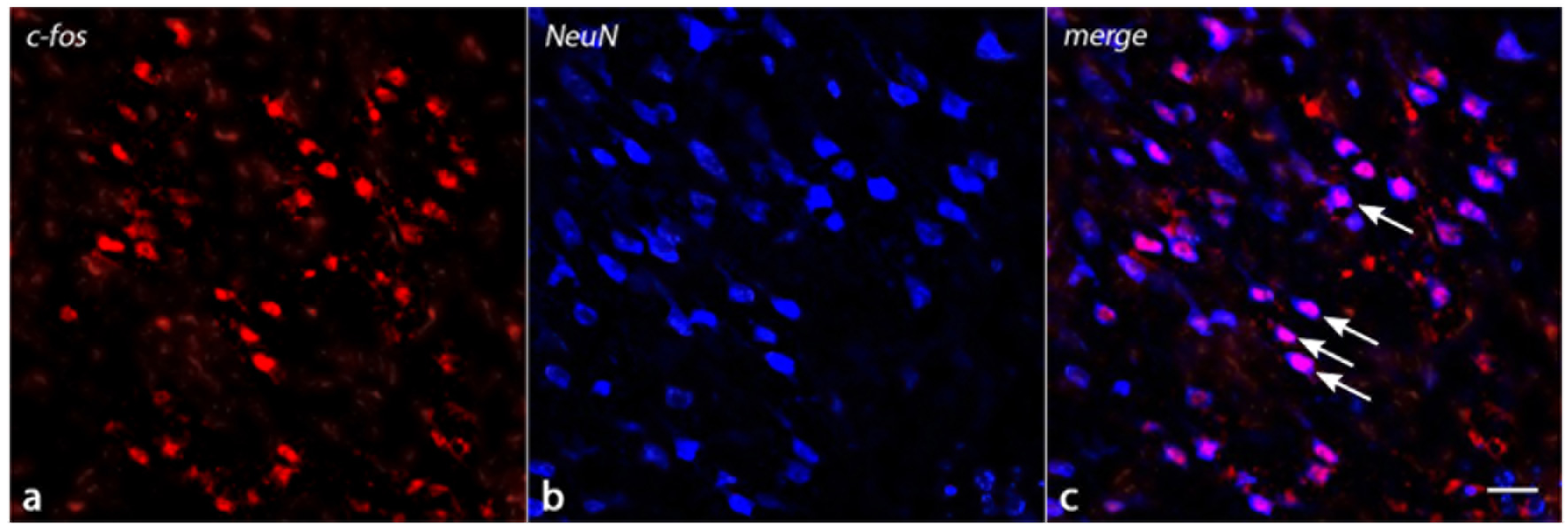
The merged image shows the colocalization of *c-fos*-labeled nuclei (red in a) to NeuN (blue in b) indicating that the induced *c-fos* activity is situated in the neuron (white arrows in c). Scale bar = 25 μm.

**Fig. 3:**
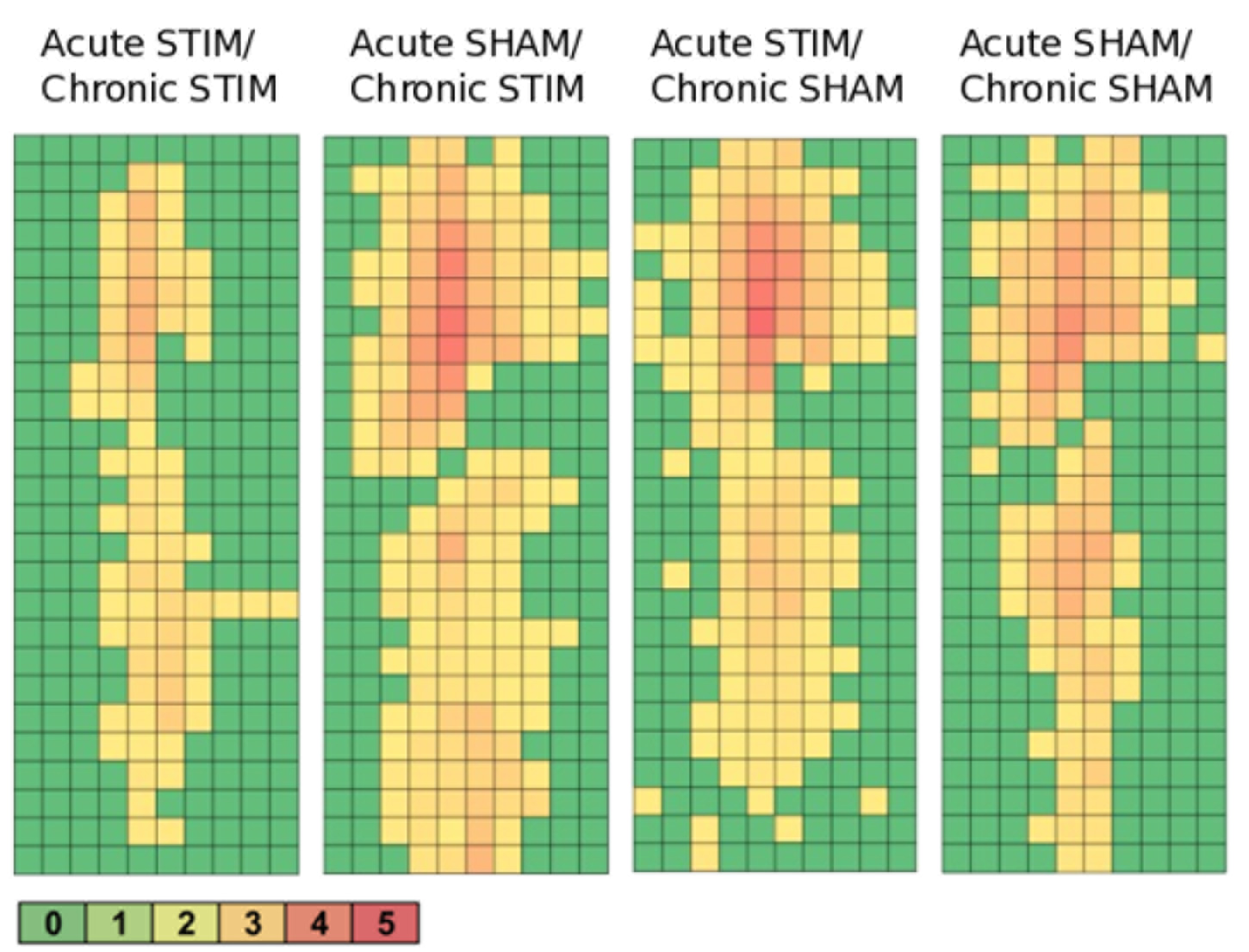
Heatmaps of *c-fos*/NeuN+ cells. Group 1 (Acute STIM/Chronic STIM) n = 4, Group 2 (Acute SHAM/Chronic STIM) n = 3, Group 3 (Acute STIM/Chronic SHAM) n = 5, Group 4 (Acute SHAM/Chronic SHAM) n = 3. Lookup-table showing number of *c-fos*/NeuN+ cells per box (100μm × 100μm).

### Effects of chronic high frequency stimulation of the dorsolateral striatum on *c-fos* expression

Chronic high frequency stimulation of the dorsolateral striatum did not change the neuronal *c-fos* expression in the region of 100 μm from the microelectrode. Statistical analysis of neurons expressing *c-fos* showed no significant reductions or increases in the number of *c-fos*/NeuN+ cells in ipsilateral striatal (p = 0.99) and cortical (p = 0.12) areas. But Group 1 showed the lowest number of *c-fos*/NeuN+ cells in comparison to the other 3 groups (Fig. 4).

**Fig. 4:**
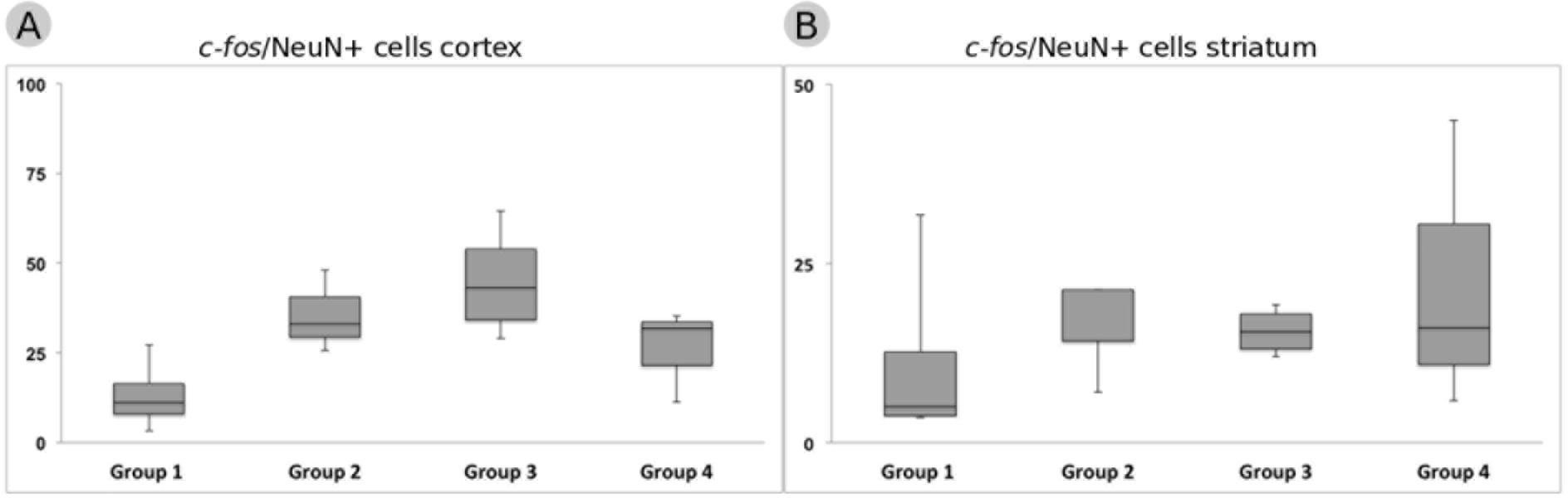
Response of *c-fos* immunoreactivity to chronic high frequency stimulation of the dorsolateral striatum. **(A)** *c-fos*/NeuN+ cells in cortical area in a distance of 100 μm from the microelectrode. **(B)** *c-fos*/NeuN+ cells in striatal area in a distance of 100 μm from the microelectrode. Whiskers showing maximum and minimum.

### Effects of the chronic implantation of a microelectrode on NeuN expression

All groups showed NeuN positive cells. Particularly strong NeuN+ cells are situated directly alongside the trajectory and represented in high density in the cortical areas in comparison to the striatum (Fig. 5). Statistical analysis of NeuN+ cells in the region of 0-100 μm from the microelectrode, in comparison to region of 400-500 μm, showed for group 1 a 95% CI of [0.37;1.16], group 2 [0.14;0.93], group 3 [1.27;2.56] und group 4 [0.37;0.95]. Thus in all 4 groups the number of NeuN+ cells in the vicinity of the microelectrode is not significantly reduced (p < 0.05). We conclude, that the chronic implantation of our stiff microelectrode does not lead to a severe neuronal cell loss and the development of a visible ‘kill zone’.

**Fig. 5:**
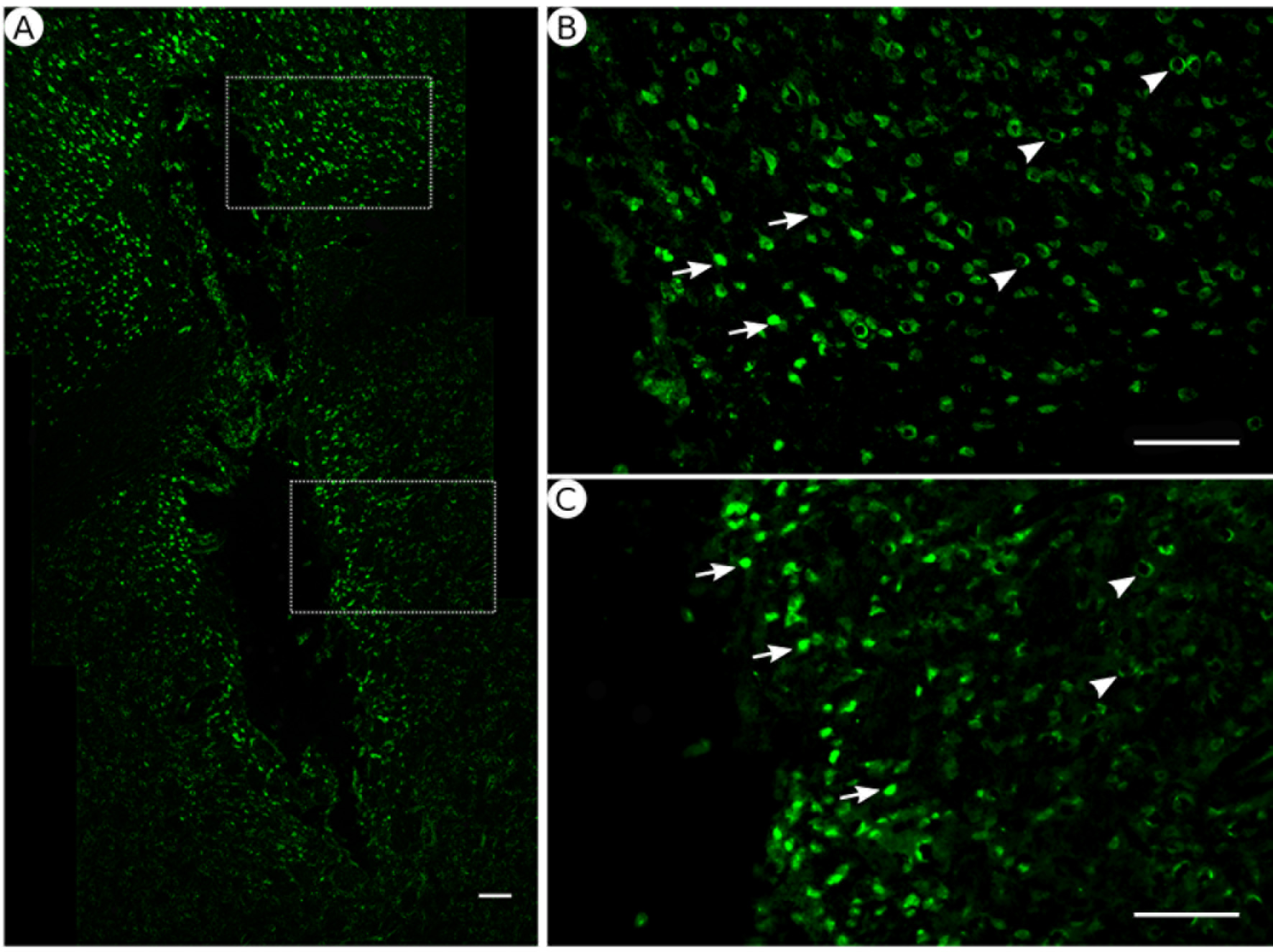
NeuN immunoreactivity. **(A)** Overview of corticostriatal area as composite of a series of coronal sections. **(B)** and **(C)** Corresponding magnified pictures from (A), white arrows pointing at high NeuN immunoreactivity, white arrowhead pointing at low NeuN immunoreactivity. Scale bar = 100 μm.

### Effects of the chronic implantation of a microelectrode on GFAP and ED1 immunoreactivity

All groups expressed glial fibrillary acidic protein (GFAP) alongside the former trajectory of the microelectrode as illustrated in Fig. 6. Astrocytes agglomerated and built a dense glial layer proximal to the implant trajectory while their typical star shape can be observed further away (Fig. 6). In Fig. 7 A, the mean background-corrected fluorescence intensities are illustrated as a function of distance to the microelectrode. The highest GFAP-immunoreactivity is found in a distance up to 100 μm of the microelectrode (peak background-corrected fluorescence intensity) and decreases with increasing distance from the trajectory. The calculation of the FWHM revealed a mean scar thickness from all groups of 129 ± 10 μm, whereas group 1 had the thickest (150 μm) and group 3 the thinnest (113 μm) FWHM (Fig. 7 B). Statistical analysis of the FWHM showed no significant difference between groups (p = 0.8826).

The chronic implantation of a stiffened polyimide microelectrode leads to a reactive astrocytosis with the formation of a glial scar with an extent of about 130 μm.

**Fig. 6:**
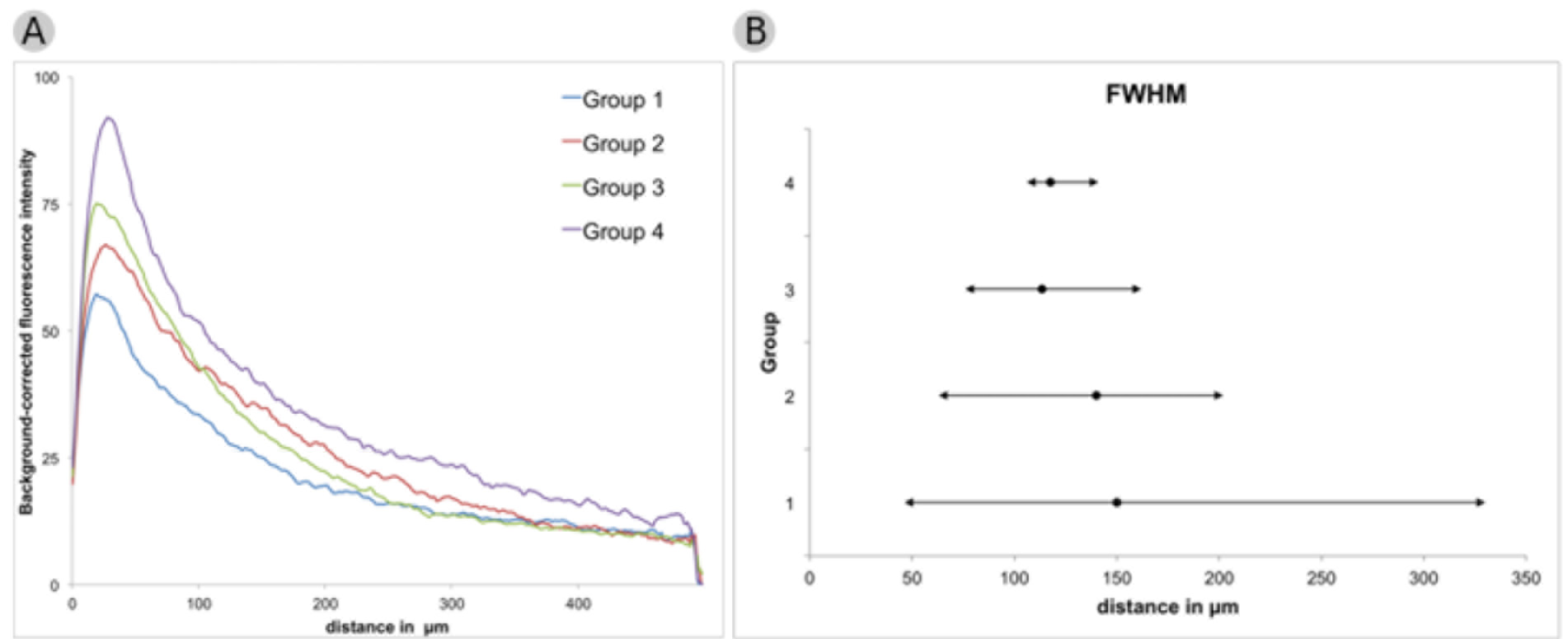
GFAP immunoreactivity. **(A)** Overview of corticostriatal area as composite of a series of coronal sections, arrowheads pointing at dense glial layer at the brain tissue/microelectrode interface. Scale bar = 100 μm. **(B)** Corresponding magnified picture from (A), white arrows pointing at GFAP+ star shaped cells (astrocytes). Scale bar = 100 μm. **(C)** Corresponding magnified picture from (B), white arrows pointing at GFAP+, star shaped cells (astrocytes). Scale bar = 25 μm.

**Fig. 7:**
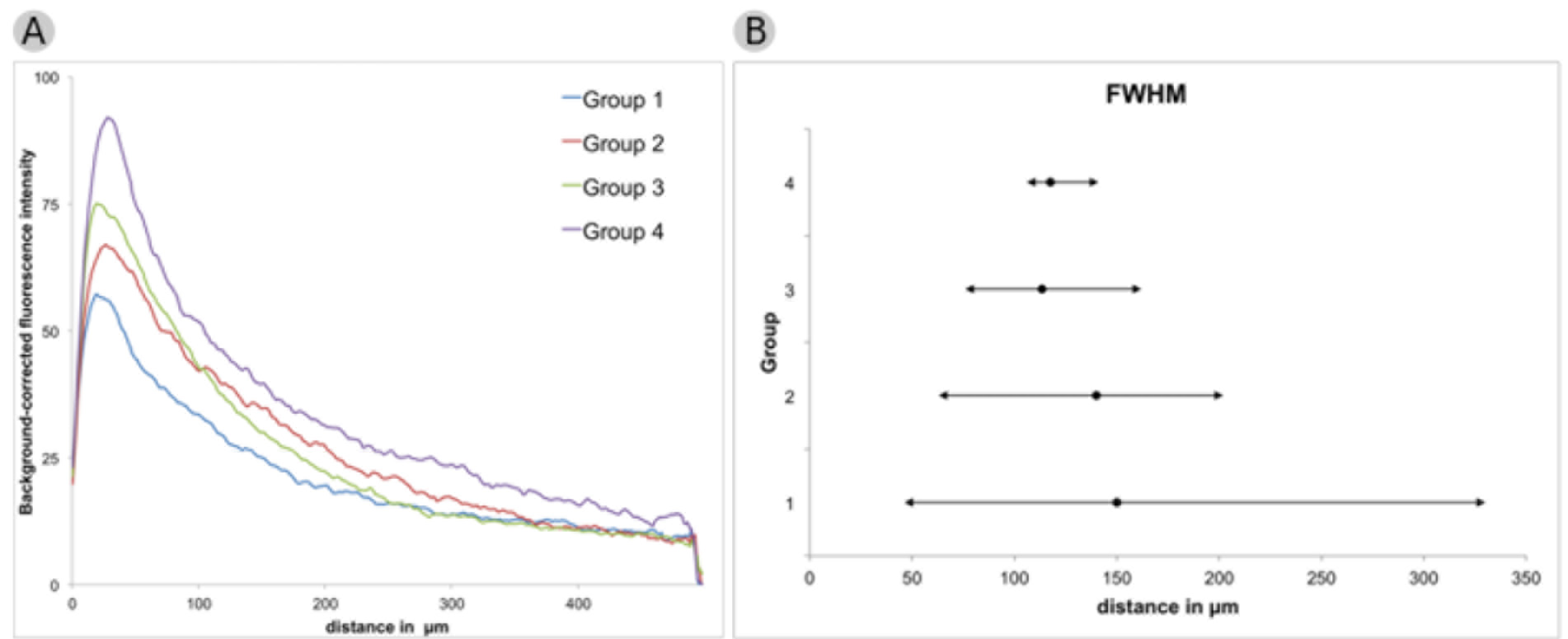
GFAP immunoreactivity. **(A)** Background-corrected GFAP fluorescence intensities illustrated as a function of distance to the microelectrode. **(B)** Full widths at half maximum of groups 1 to 4. Mean + maximum/minimum

Due to the fact that we implanted an otherwise flexible microelectrode glued to a glass rod, we also analyzed the GFAP immunoreactivity in dependence of the microelectrode orientation. Fig. 8 illustrates the background-corrected fluorescence intensities separated in medial and lateral, in dependence of the microelectrode orientation (lateral (A), medial (B)). The line graphs show that there is no difference between the lateral and medial GFAP expression. The astrocytic reaction seems to be independent from the utilized material.

**Fig. 8:**
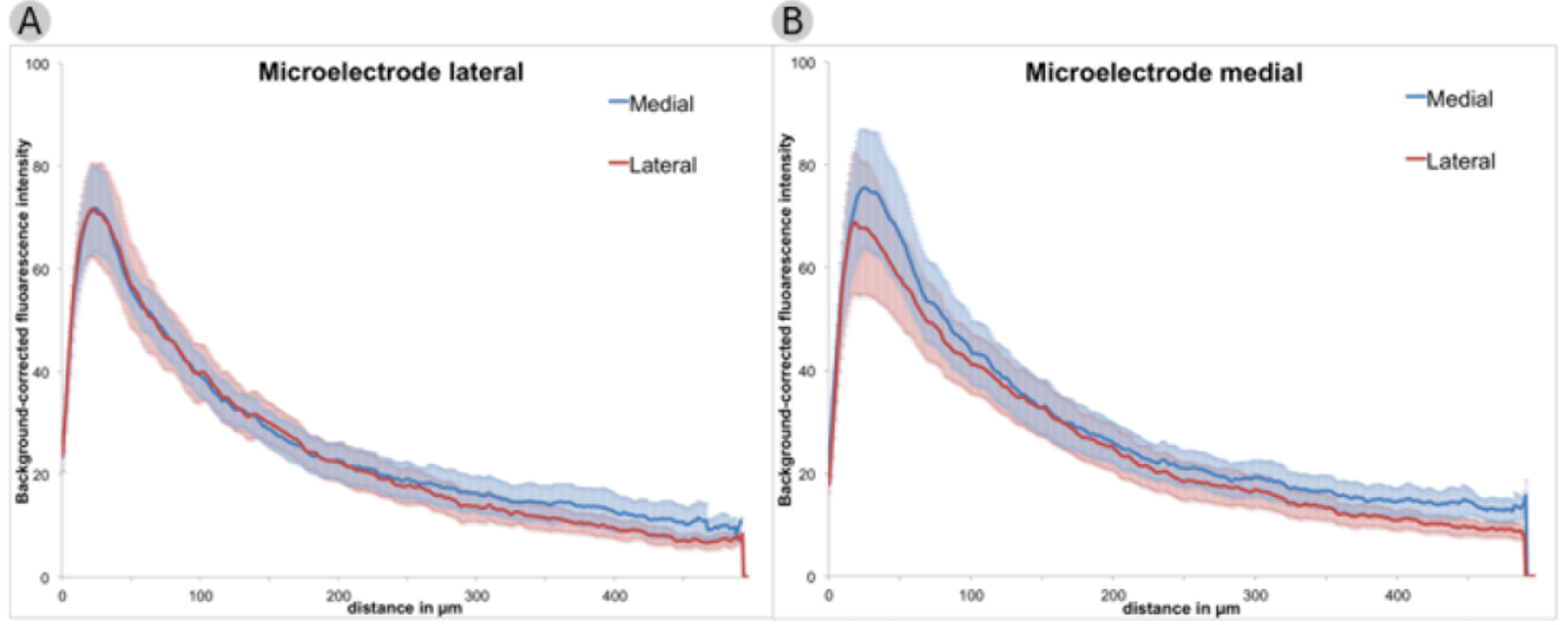
Background-corrected GFAP immunoreactivity in dependence of the microelectrode direction. **(A)** Microelectrode direction lateral, Background-corrected GFAP immunoreactivities separated in medial and lateral. **(B)** Microelectrode direction medial, Background-corrected GFAP immunoreactivities separated in medial and lateral. Means ± SE.

ED1 expression was present, but barely visible along the whole trajectory. An exemplary picture is shown in Fig. 9. ED1 positive cells formed agglomerates and could hardly be distinguished from each other.

**Fig. 9:**
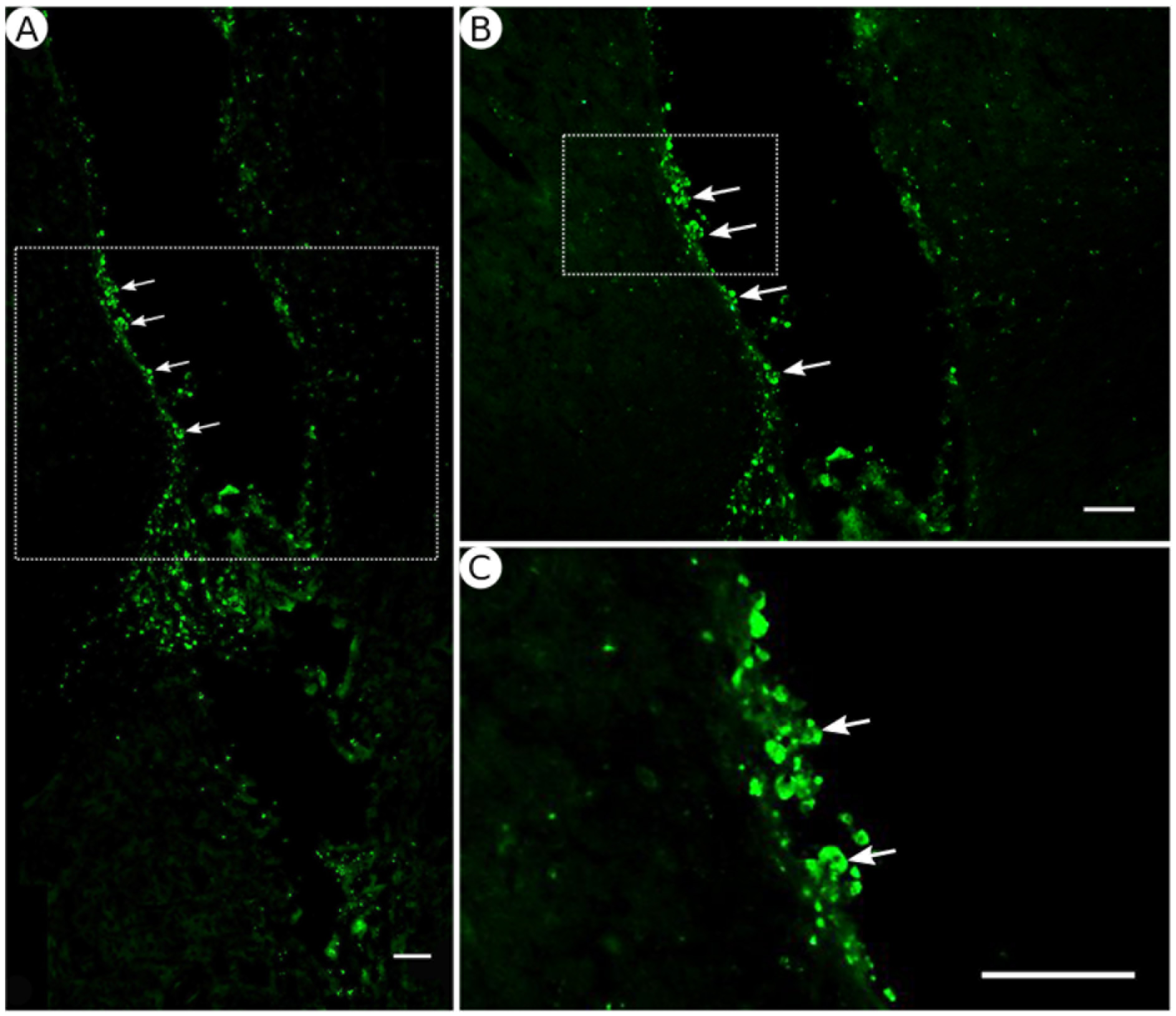
ED1 immunoreactivity. **(A)** Overview of corticostriatal area as composite of a series of coronal sections, arrows pointing at ED1+ cells at the brain tissue/microelectrode interface. Scale bar = 100 μm. **(B)** Corresponding magnified picture from (A), white arrows pointing at ED1+ cells (microglia). Scale bar = 100 μm. **(C)** Corresponding magnified picture from (B), white arrows pointing at agglomerated ED1+ cells in the lumen. Scale bar = 25 μm.

## DISCUSSION

The *c-fos* immunochemistry is widely used as marker for neuronal activation (Faull, 1989, Bullitt, 1990, Herrera and Robertson, 1996, Wilson et al., 2002) despite the fact that *c-fos* expression is relatively unspecific and can be induced by chemical, physical or electrical stress, and has even been constitutively expressed in certain brain areas (Greenberg and Ziff, 1984, Morgan and Curran, 1989, Hughes et al., 1992, Herrera and Robertson, 1996). However in our study, the highest density of activated neurons is located directly alongside the trajectory, and declines with increasing distance from the implantation region (Fig. 3). This observation indicates that the *c-fos* expression is caused by the implanted microelectrode, and is reinforced by the finding of a low basal *c-fos* expression in the contralateral brain area. Statistical analysis verified the significantly elevated number of *c-fos*/NeuN positive cells in the vicinity of the microelectrode. As such, the implantation of a microelectrode can lead to an adaptation of the surrounding neurons, as expressed through a change in their gene expression and translation. One possible explanation could be that the CNS injury caused by the implantation of the microelectrode leads to a release of cytokines and growth factors, resulting in a modified cellular state of action (Polikov et al., 2005, He et al., 2007, Pineau and Lacroix, 2007). The cells involved in the foreign body reaction, especially microglia and astrocytes, change in response to the trauma their state of action and release cytokines, growth factors, enzymes and other neuroactive substances (Eddleston and Mucke, 1993, Fawcett and Asher, 1999, Loane and Byrnes, 2010, Benarroch, 2013). Furthermore, Herrera and Robertson and others showed that cortical brain injury can cause an elevated *c-fos* expression in the surrounding neurons as a response to the release of excitatory amino acids (Faden et al., 1989, Herrera and Robertson, 1990, Sharp et al., 1990). Recent studies utilizing Fast Cyclic Voltammetry (FCV) during human DBS implantations has demonstrated this “microthalamotomy” dubbed release of adenosine (Bennet et al., 2016).

Chronic high frequency stimulation of the dorsolateral striatum did not significantly change the number of *c-fos*/NeuN positive cells in the stimulated area along the lower end of the trajectory (Fig. 1) As the chosen stimulation parameters (60μs) coincide with known chronaxy values for neuronal fibres, but not somata, we would rather expect an axonal than somal stimulation (Holsheimer et al., 2000, Jan Holsheimer, 2000, McIntyre et al., 2004, Löffler and Lujàn, 2016). So one would not expect a local *c-fos* expression due to electrical HFS, but rather a change in afferent and efferent neuronal cells. Considering the broad somatotopically organized cortico-striatal connections (Voorn et al., 2004), we looked at the *c-fos* expression of cortical neurons. In comparison to the other groups, Group 1 (Acute STIM/Chronic STIM) had shown the lowest number of *c-fos*/NeuN positive cells. This effect could be a result of an antidromic inhibition by cortico-striatal pathways. HFS of the dorsolateral striatum could lead through an antidromic axonal stimulation to a suppression of the activity of cortical neurons, i.e. lower *c-fos* expression, following the GABAergic mode of action, through facilitatory GABA autoreceptors (Li et al., 2007, Feuerstein et al., 2011). This is keeping in mind that GABA acts as an inhibitory transmitter and, *c-fos* expression follows excitatory amino acid release (Faden et al., 1989, Hughes and Dragunow, 1995).

The chronic implantation of a polyimide microelectrode leads to the formation of a glial scar and accumulation of microglia cells at the electrode-brain interface. A significant neuronal cell loss in the vicinity of the microelectrode was not observed.

In our study, we used polyimide microelectrodes with a 125 μm glas rod glued to their back, to achieve precisely located electrical microstimulation in the brain (Fig. 1). After a mean implantation time of 70 days we detected an accumulation of astrocytes (high GFAP expression) alongside the implanted microelectrode trajectory. The highest GFAP intensities are found within 100 μm from the microelectrode with a mean glial scar width of 130 μm, whereas the density of GFAP+ cells decreases as the distance to the microelectrode increases (Fig. 7). The GFAP expression gave no indication for a material-dependent astrocytic reaction, as fluorescence distribution was similarly medial and lateral, resp. on PI or glass (Fig. 8). Astrocytes represent 30-65% of the glial cell population in the CNS, and are essential for maintaining a proper neuronal environment (Nathaniel and Nathaniel, 1981, Kimelberg et al., 1993). Following chronic implantation of an electrode, astrocytes are thought to build an encapsulation (glial scar) (Turner et al., 1999). Our findings are in accordance with reports suggesting a formation of an astrocytic boundary around the lesion, building up a tightly connected network of hyperfilamentuos scar astrocytes, surrounded by extracellular matrix (Bush et al., 1999, Fawcett and Asher, 1999, Faulkner et al., 2004, Seifert et al., 2006).

The ED1 immunoreactivity as a representative of activated monocytes and macrophages is weak. Monocytes and macrophages are one of the first responders after injury or lesion of the CNS and react to the change by activation (Kreutzberg, 1996). Activated monocytes begin to proliferate and change their morphology to a more “amoeboid” shape (Polikov et al., 2005). We also observed this with residuals of the ED1+ cells (Fig. 9). These cells were particularly notable at the electrode-brain interface. Over time, the acute activity and thus cellular foreign body reaction declines as expected by a mean implantation time of 10 weeks (Potter et al., 2012).

The vitality of the surrounding neurons is essential for a stable signal transmission between electrode and CNS (Biran et al., 2005). The foreign body reaction, following electrode implantation, can lead to neuronal cell loss and the formation of a ‘kill zone’ (Edell et al., 1992, McConnell et al., 2009b). The extent of this ‘kill zone’ can reach up to 100 μm (Polikov et al., 2005). In our study, we could not detect a significant neuronal cell loss in the vicinity of the chronically implanted microelectrode. This could be due to the weak foreign body reaction with a low ED1 expression, since neuronal degeneration correlates with local inflammation (McConnell et al., 2009b). In studies that compared the implantation of a electrode with the sole stab injury with the electrode, no significant neuronal cell loss in the stab groups were found (Biran et al., 2005, McConnell et al., 2009b, Potter et al., 2012). Considering the advances in electrode designs, material and implantation method, our chronically implanted polyimide microelectrode preserved viable neurons in its immediate vicinity.

To conclude, our study has shown that sole chronic implantation of a polyimide microelectrode - and not electrical stimulation - leads to a *c-fos* expression along its trajectory. The neuronal cell density in immediate vicinity to the microelectrode was not significantly reduced, presumably due to the low local inflammation.

